# Zinc finger RNA binding protein Zn72D regulates ADAR-mediated RNA editing in neurons

**DOI:** 10.1101/631986

**Authors:** Anne L. Sapiro, Emily C. Freund, Lucas Restrepo, Huan-Huan Qiao, Amruta Bhate, Qin Li, Jian-Quan Ni, Timothy J. Mosca, Jin Billy Li

## Abstract

Adenosine-to-inosine RNA editing, catalyzed by ADAR enzymes, alters RNA sequences from those encoded by DNA. These editing events are dynamically regulated, but few *trans* regulators of ADARs are known *in vivo*. Here, we screen RNA binding proteins for roles in editing regulation using *in vivo* knockdown experiments in the *Drosophila* brain. We identify Zinc-Finger Protein at 72D (Zn72D) as a regulator of editing levels at a majority of editing sites in the brain. Zn72D both regulates ADAR protein levels and interacts with ADAR in an RNA-dependent fashion, and similar to ADAR, Zn72D is necessary to maintain proper neuromuscular junction architecture and motility in the fly. Furthermore, the mammalian homolog of Zn72D, Zfr, regulates editing in mouse primary neurons, demonstrating the conservation of this regulatory role. The broad and conserved regulation of ADAR editing by Zn72D in neurons represents a novel mechanism by which critically important editing events are sustained.

## Introduction

RNA editing expands genetic diversity by altering bases encoded by the genome at the RNA level (Eisenberg and Levanon, 2018; Nishikura, 2016; Walkley and J. B. Li, 2017). The deamination of adenosine (A) into inosine (I), a highly prevalent form of mRNA editing, is catalyzed by adenosine deaminase acting on RNA (ADAR) proteins, which are double-stranded RNA binding proteins that are conserved in metazoans (Bass, 2002). Inosine is recognized by the cellular machinery as guanosine (G); therefore, a single editing event in an RNA has the ability to change the regulation of the RNA or to change the protein encoded by the transcript by altering a codon or splice site (Nishikura, 2010). Millions of these RNA editing sites have been identified, necessitating a better understanding of how this process is regulated (Walkley and J. B. Li, 2017).

Proper regulation of ADAR proteins and A-to-I RNA editing is essential to organismal health. Humans have two catalytically active ADAR proteins, and functional changes in both proteins are associated with disease. ADAR1 edits endogenous double-stranded RNA, which is critical for proper innate immune function (Liddicoat et al., 2015; Mannion et al., 2014; Pestal et al., 2015), and loss of ADAR1 sensitizes tumors to regression (Gannon et al., 2018; Ishizuka et al., 2018; Liu et al., 2018). ADAR2 regulates the editing of a number of ion channels important for regulating neuronal excitability (Rosenthal and Seeburg, 2012), and its dysregulation is associated with a host of neurological diseases including amyotrophic lateral sclerosis, astrocytoma, and transient forebrain ischemia (Slotkin and Nishikura, 2013). In *Drosophila*, loss of the single *Adar* homologue, most akin to mammalian *Adar2*, leads to neurological phenotypes including impaired locomotion and age-related neurodegeneration (Palladino et al., 2000). While maintaining RNA editing levels is critical for proper immune and neuronal function, regulation of ADAR proteins and editing levels is poorly understood.

Recent studies suggest that regulation of RNA editing levels is highly complex and that critical RNA editing regulators are yet to be identified. RNA editing levels differ across tissues and developmental stages, and these changes do not always correlate with *Adar* mRNA or protein expression (J. B. Li and Church, 2013; Sapiro et al., 2019; Tan et al., 2017; Wahlstedt et al., 2009). *Trans* regulators of ADAR proteins may help explain this variation in editing levels (Sapiro et al., 2015); however, few ADAR and editing level regulators are known. In mammals, Pin1, WWP2, and AIMP2 regulate ADAR protein levels or localization, which can then alter editing levels (Behm et al., 2017; Marcucci et al., 2011; Tan et al., 2017). Editing regulators can also be site-specific, meaning they regulate ADAR editing at only a subset of editing sites rather than globally regulating ADAR activity. Studies in *Drosophila* identified FMR1 and Maleless as site-specific regulators of editing (Bhogal et al., 2011; Reenan et al., 2000). Further study has verified that human homologs of both FMR1 (Tran et al., 2019) and Maleless (Hong et al., 2018a), along with a number of other RNA binding proteins and splicing factors, act as site-specific regulators of RNA editing. These factors, including SRSF9, DDX15, TDP-43, DROSHA, and Ro60 (Garncarz et al., 2013; Quinones-Valdez et al., 2019; Shanmugam et al., 2018; Tariq et al., 2013), help to explain some variation in editing levels; however, with thousands of editing sites in flies and millions in humans (Ramaswami and J. B. Li, 2014), additional regulators likely remain undiscovered. These previous studies highlight RNA binding proteins as strong candidates for editing regulators (Washburn and Hundley, 2016). Because of the conserved roles of *Drosophila* editing regulators as well as the ability to measure nervous system phenotypes, flies serve as an important model for understanding the regulation of editing as it relates to human neurological diseases.

To identify novel regulators of RNA editing in the brain, we screened 48 RNA binding proteins for regulation of editing levels using RNA-interference (RNAi) in *Drosophila* neurons. We identified *Zinc-Finger Protein at 72D* (*Zn72D*) as a novel regulator of RNA editing as *Zn72D* knockdown altered editing at nearly two-thirds of assayed editing sites. *Zn72D* knockdown led to a decrease in ADAR protein levels, although this decrease did not fully explain the editing level changes. We further determined that Zn72D and ADAR physically interact in the brain by binding the same RNA species. In addition to editing changes, loss of Zn72D also led to defects at the neuromuscular junction (NMJ) and impaired locomotion in the fly. Finally, we found that the mouse homolog of Zn72D, Zfr, regulates editing levels in primary neuron cultures, suggesting this mode of editing regulation is highly conserved.

## Results

### An RNAi screen identifies Zn72D as a novel regulator of RNA editing

To better understand how ADAR editing is regulated in the brain, we designed an *in vivo* screen to identify novel regulators of editing in *Drosophila*. Since RNA binding proteins (RBPs) play critical roles in RNA processing events and regulate a number of editing events in flies and mammals (Washburn and Hundley, 2016), we chose to focus on RBPs as candidate regulators of editing. We created a collection of UAS-shRNA lines targeting annotated RBPs as well as Green Fluorescent Protein (GFP) as a control, as done previously (Ni et al., 2011). To assay whether loss of these RBPs influenced editing levels, we designed a simple screen (**Figure 1A**), by crossing UAS-shRNA lines targeting an RBP or GFP to the pan-neuronal driver *C155-Gal4*. We then extracted RNA, produced, and sequenced RNA-sequencing (RNA-seq) libraries from two biological replicates of adult knockdown brains. We determined editing level differences at known editing sites between control and knockdown brains. To validate this approach, we first checked the reproducibility of editing levels between biological replicates of GFP RNAi brains used in the screen as a control and found that editing levels between replicates were highly reproducible (**Figure 1B**). We then tested the design of the screen by knocking down *Adar* using two independent shRNA lines (BDSC28311 and VDRC7763), which reduced *Adar* mRNA levels by 60% and 72%, respectively. We compared editing levels between two replicates of each *Adar* knockdown and their matched replicates of GFP knockdown at previously identified editing sites. To avoid looking at potential SNPs or false positive editing sites, we limited the sites queried in our screen to high-confidence editing sites – those that were reproducibly edited and altered by *Adar* knockdowns in these pilot experiments. In total, we identified 1236 editing sites that were reproducibly edited between the independent sets of GFP knockdown replicates and were reduced significantly by the stronger *Adar* knockdown as measured by Fisher’s exact tests (**Figure 1C**).

**Figure 1.**
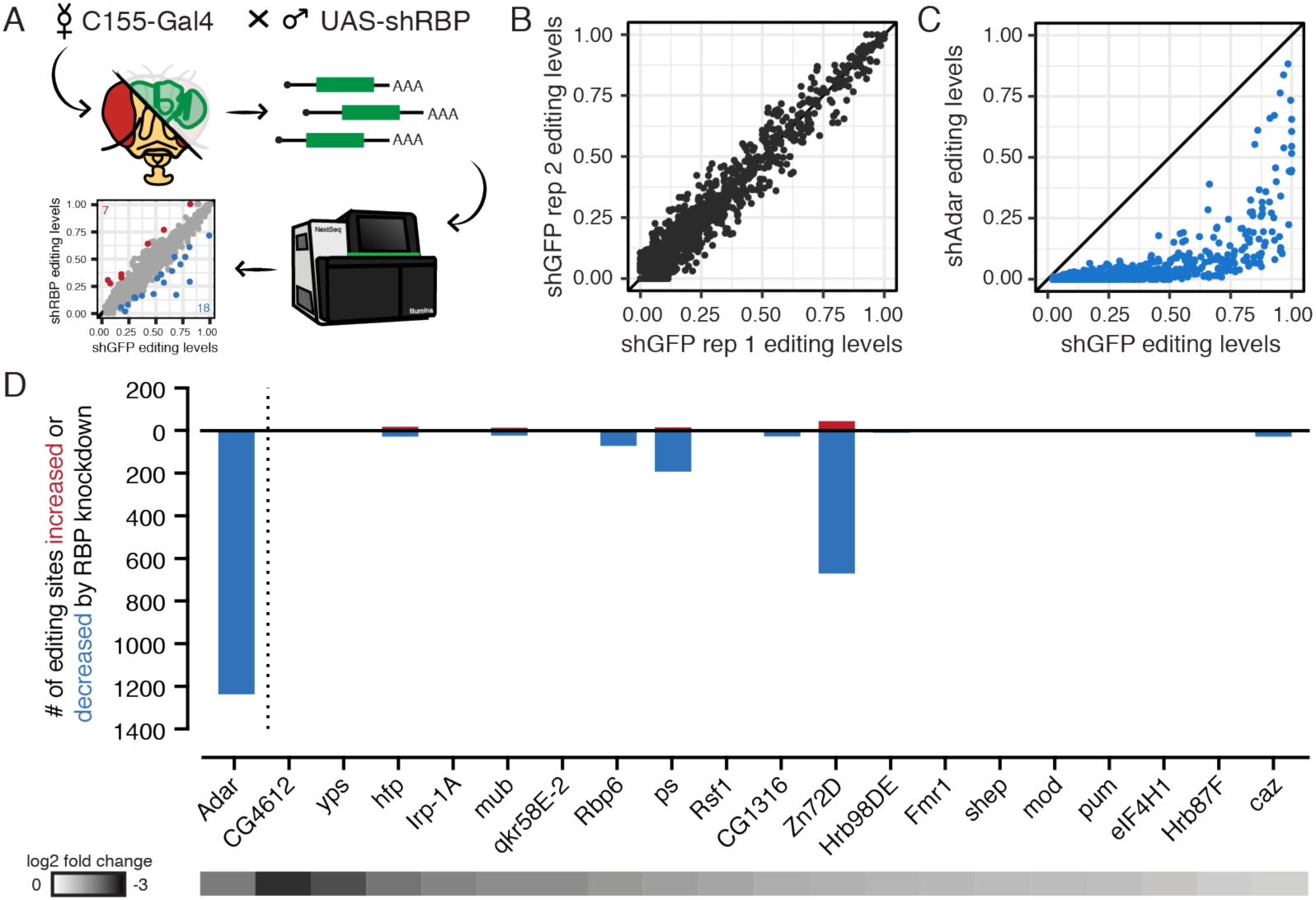
An RNAi screen identifies Zn72D as a novel regulator of RNA editing. (A) Schematic of RNAi screen. Pan neuronal Gal4 driver *C155-Gal4* was crossed to flies containing UAS-shRNAs targeting 1 of 20 different RNA binding proteins. RNA from 3-5 day old adult females was sequenced to compare editing levels between *C155-Gal4; UAS-shGFP* controls and *C155-Gal4; UAS-shRBP* flies. (B) Comparison of editing levels across two biological replicates of *shGFP* controls. Biological replicates were highly reproducible. (C) Comparison of editing levels between *C155-Gal4; UAS-shGFP* and *C155-Gal4; UAS-shAdar* at sites used in the screen. All sites are reduced. Blue dots, *p* < 0.05, Fisher’s exact tests. (D) The number of editing sites found to be increased or decreased (*p* < 0.05, Fisher’s exact tests) upon knockdown of each of 20 RBPs screened. The heat map below shows the log_2_ fold change of each target RBP between knockdown and control as measured by mRNA levels from RNA-seq. *shZn72D* shows the greatest number of altered sites besides *shAdar*.

After validating our screening strategy, we crossed shRNA lines targeting 48 different RBPs, starting with RBPs that are highly expressed in the brain (see **Table S1**). Of the 48 knockdowns, 17 caused lethality before adulthood and were not screened for editing changes. For the 31 knockdowns that produced viable adults, we performed qPCR to determine the level of knockdown of the target. 19 RNA binding protein targets showed greater than 40% knockdown efficiency, and RNA-seq libraries from two replicates of each and two GFP-targeting control knockdown libraries were sequenced. We then determined editing levels at the 1236 sites that were affected by *Adar* knockdowns. Editing levels between all biological replicates used in the screen were highly reproducible, similar to shGFP replicates (**Figure S1A**). We determined whether sites differed between shGFP controls and RBP knockdowns using Fisher’s exact tests of total A and G counts from the biological replicates combined. **Figure 1D** shows the number of editing sites that were more highly or lowly edited in each knockdown than in the GFP controls, as well as the knockdown efficiency for each target as measured by RNA-sequencing (see **Tables S2**, **S3**). The majority of the RBP knockdowns showed evidence of positive or negative regulation of editing at fewer than 50 editing sites, and these effects generally led to small changes in editing (**Figure S1B**). Two RBP knockdowns had slightly wider-ranging effects on editing levels. Knockdown of *Rbp6* decreased editing at 72 sites and increased editing at 2 sites, and knockdown of *pasilla* decreased editing at 193 sites and increased editing at 15 sites. By far the most robust regulator of RNA editing, however – in terms of both the number of sites altered and the strength of the effect – was *Zinc-finger protein at 72D (Zn72D)*, knockdown of which decreased editing at 670 editing sites and increased editing at 44 sites, affecting 59% of the sites measured. This dramatic regulation of editing greatly exceeded that of all other RBPs screened as well as many other RBPs reported to regulate ADAR editing (Washburn and Hundley, 2016); therefore, we focused on characterizing Zn72D in this work.

### *Zn72D* knockdown alters editing levels at a distinct subset of editing sites

As Zn72D was the strongest hit to come out of our screen for editing regulators, we took a closer look at the sites that were affected by *Zn72D* knockdown. Comparing editing levels between *C155-Gal4; UAS-shGFP* and *C155-Gal4; UAS-shZn72D* revealed dramatic changes in editing at many, but not all, sites (**Figure 2A**), suggesting Zn72D is a site-specific editing regulator. To validate this striking editing phenotype, we crossed an independent *UAS-shZn72D* line (obtained from Bloomington Drosophila Stock Center) to *C155-Gal4* and sequenced the RNA to confirm the editing level differences from control knockdown. We observed a similar editing phenotype with this independent shRNA line, with the same editing sites showing the same responsiveness to *Zn72D* knockdown with both shRNAs (**Figure S2A-B, Table S2**). To verify that the editing phenotype was not a consequence of the RNAi system itself or off target effects, we crossed two *Zn72D* mutant alleles, *Zn72D^1^* and *Zn72D^1A14^*, which caused premature stop codons at amino acids 38 and 559 respectively. These *Zn72D^1/1A14^* mutants died before reaching adulthood, as previously reported (Brumby et al., 2004), so we collected heads from pupae approximately 72 hours after puparium formation and sequenced the RNA to check editing levels. *Zn72D* mutant pupal heads also showed large differences in editing from wild type pupal heads (**Figure 2B, Table S2**). We then compared the changes in editing observed in the *Zn72D* knockdowns to those in the *Zn72D* mutants. Despite the difference between the developmental stages of the flies, the editing level differences between wild type and *Zn72D* mutant pupal heads were similar to the editing level differences found at the same sites between shGFP and shZn72D in adult brains (**Figure 2C**), confirming that the *Zn72D* editing phenotype was highly reproducible and site-specific.

**Figure 2.**
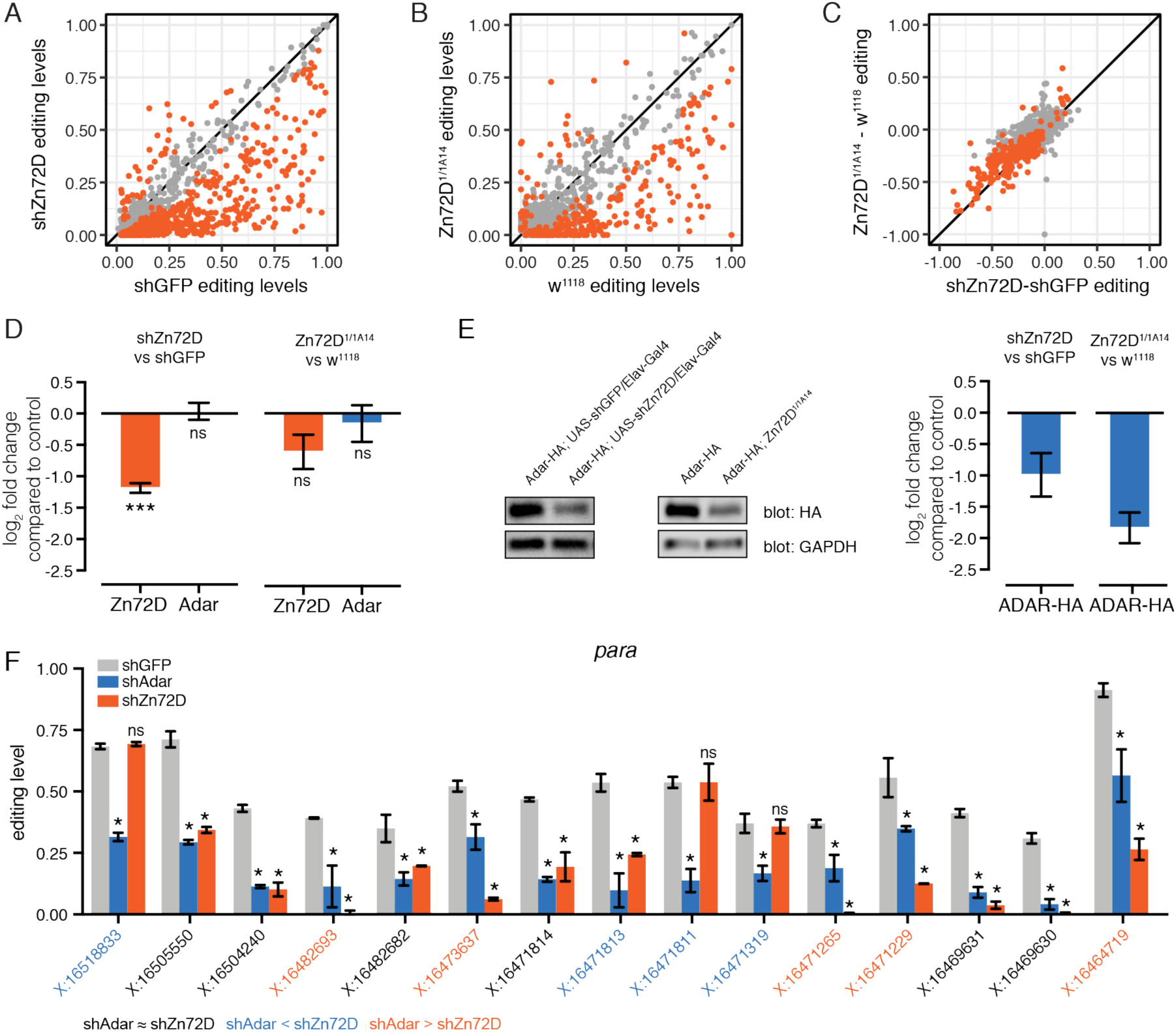
Zn72D regulates ADAR editing at a distinct subset of editing sites. (A) Comparison of editing levels at individual editing sites (dots) between *C155-Gal4; UAS-shGFP* and *C155-Gal4; UAS-shZn72D*, from the RNAi screen in Figure 1. Orange dots, *p* < 0.05, Fisher’s exact tests. (B) Comparison of editing levels between *w^1118^* and *Zn72D^1/1A14^* pupal heads. Orange dots, *p* < 0.05, Fisher’s exact tests. The majority of sites are altered in both *Zn72D* knockdowns and mutants compared to controls. (C) Comparison of the difference in editing between *C155-Gal4; UAS-shZn72D* and *C155-Gal4; UAS-shGFP* and *Zn72D^1/1A14^* and *w^1118^* from (A) and (B). The same sites are significantly altered in both knockdown and mutants. Orange dots, *p* < 0.05 in both. (D) Log_2_ fold change of *Zn72D* and *Adar* mRNA levels in *C155-Gal4; UAS-shZn72D* compared to *C155-Gal4; UAS-shGFP* adult brains and *Zn72D^1/1A14^* compared to *w^1118^* pupal heads. *Adar* mRNA levels are not decreased in *Zn72D* knockdown and mutants. ***, *p* < 0.0001, ns, *p* > 0.05, Wald tests. (E) Western blot of ADAR-HA protein in *Elav-Gal4/shGFP* and *Elav-Gal4/shZn72D* adult brains and *w^1118^* and *Zn72D^1/1A14^* pupal heads. At right, quantification of HA loss in *Zn72D* knockdown and mutant compared to controls, normalized to GAPDH. ADAR-HA protein levels are decreased in both *Zn72D* knockdown and mutants. n=3 for each comparison. (F) Editing levels in *C155-Gal4; UAS-shGFP*, *C155-Gal4; UAS-shAdar*, and *C155-Gal4; UAS-shZn72D* brains in *para*. Sites within the transcript are differentially affected by *Zn72D* loss. Bars represent mean, error bars represent +/− SD (n=2). *, *p* < 0.001, ns, *p* > 0.05, between *shGFP* and either *shAdar* (above blue bar) or *shZn72D* (above orange bar), Fisher’s exact tests. Orange coordinates, shAdar editing less than shZn72D editing. Black coordinates, no difference between *shAdar* and *shZn72D*. Blue coordinates, shAdar editing less than shZn72D editing. Blue and orange, *p* < 0.001, Fisher’s exact tests.

Since 93% of the editing sites affected by *Zn72D* knockdown had decreased editing levels, we wanted to determine whether Zn72D loss reduced *Adar* mRNA or protein levels. We first checked *Adar* mRNA levels between *shGFP* and *shZn72D* brains and wild type and mutant heads. Decrease of *Zn72D* did not lead to a significant decrease in *Adar* mRNA levels (**Figure 2D, S2C**). To check ADAR protein levels, we knocked down *Zn72D* in the *Adar-HA* (Jepson et al., 2011) background, in which endogenous ADAR protein is tagged with HA, using the pan neuronal driver *Elav-Gal4*. By western blot, we found that ADAR-HA protein was decreased by 49% in *Zn72D* knockdown brains. We also crossed the mutants into the *Adar-HA* background, and we found ADAR-HA levels were decreased by 72% in *Adar-HA; Zn72D^1/1A14^* pupal heads (**Figure 2E**), verifying an ADAR protein reduction upon loss of Zn72D.

While a decrease in ADAR protein may explain some editing decreases in *Zn72D* knockdown and mutants, this finding could not fully explain the complex editing phenotype observed. The editing phenotype in *Zn72D* knockdown clearly differed from both the strong *Adar* knockdown (see **Figure 1C**) and the weaker *Adar* knockdown (see **Figure S3A**) used to validate our screening approach, which caused a global decrease in editing at all sites. Unlike the *Adar* knockdowns, the two *Zn72D* knockdowns affected 59% and 66% of editing sites (see **Figure 2A**, **S2A**). The sites that are unaffected are not influenced by the global decrease of ADAR protein level. We further examined the editing phenotype of sites located within the same transcript. Of 187 transcripts where we looked at multiple editing sites, 131 (70%) included at least one site that was affected and at least one site that was not affected by Zn72D knockdown, and the vast majority of transcripts with many editing sites showed mixed effects within transcripts (**Figure S3B-C**). For example, the highly edited transcript *paralytic (para)* had multiple editing sites that showed differential editing changes in response to *Zn72D* knockdown. **Figure 2F** shows editing levels at 15 highly edited sites (>20% in controls) in *para* in *shGFP*, *shAdar* (60% knockdown) and *shZn72D* brains. At 6 sites, *Zn72D* and *Adar* knockdowns led to similar editing decreases (Figure 2F, highlighted in black), whereas at 4 sites *Adar* knockdown decreased editing more than *Zn72D* knockdown (Figure 2F, highlighted in blue), and at 5 sites *Zn72D* knockdown decreased editing more than *Adar* knockdown (Figure 2F, highlighted in orange). These sites with different responses to *Zn72D* knockdown could be found within a few bases of each other, as seen at three *para* editing sites located within 4 bases of each other (chrX:16471811 to chrX:16471814). Another transcript, *quiver* (*qvr)*, showed similar patterns, including vastly different effects of *Zn72D* knockdown on four sites that were all more than 70% edited in controls and located within 23 bases of each other (chr2R:11447601 to chr2R:11447623) (**Figure S3D**). These results suggested that Zn72D’s site-specific effect on editing was highly localized down to individual editing sites, which is not what we would expect if Zn72D simply regulated ADAR protein levels.

### Zn72D interacts with ADAR and binds ADAR-target mRNAs

Since decreases in ADAR protein levels did not explain the Zn72D knockdown editing phenotype, we hypothesized that site-specific regulation of editing by Zn72D might result from the protein binding the same transcripts ADAR binds, as has been previously demonstrated for other known site-specific regulators of editing (Bhogal et al., 2011; Hong et al., 2018; Quinones-Valdez et al., 2019; Rajendren et al., 2018; Shanmugam et al., 2018). We first asked whether Zn72D and ADAR proteins were both found in the nucleus, where editing occurs (Rodriguez et al., 2012). Utilizing *Zn72D-GFP* flies that express a GFP-tagged version of the Zn72D from the endogenous locus (Morin et al., 2001), we used immunofluorescence microscopy to determine the localization of both ADAR and Zn72D proteins in *Adar-HA; Zn72D-GFP* flies. We found that both ADAR and Zn72D colocalize broadly to neuronal nuclei within the brain, along with a nuclear marker in neurons, Elav (**Figure 3A-D**).

**Figure 3.**
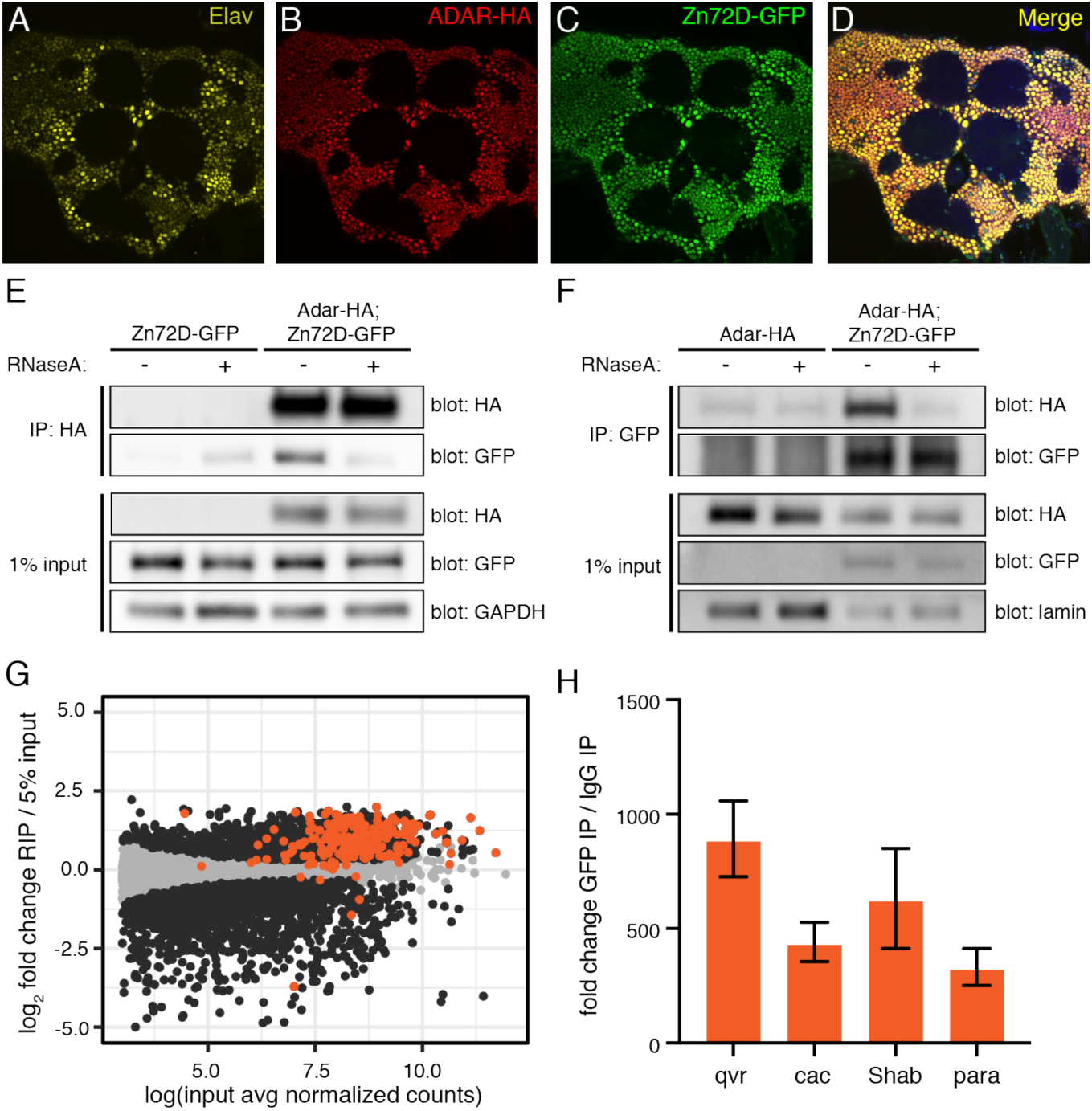
Zn72D interacts with ADAR in an RNA-dependent manner. (A-D) Immunofluorescent staining of Elav (A), ADAR-HA (B), and Zn72D-GFP (C) and all three merged (D) in the adult fly brain. All proteins are expressed in neuronal nuclei. (E) Western blots of HA and GFP following co-immunoprecipitation of ADAR-HA from *Zn72D-GFP* (control) and *Adar-HA; Zn72D-GFP* heads. Half of each IP was treated with RNase A. Blots of HA, GFP, and GAPDH from 1% of input material are shown. n=3. (F) Western blots of HA and GFP following co-immunoprecipitation of *Zn72D-GFP* from *Adar-HA; Zn72D-GFP* head nuclei. Half of each IP was treated with RNase A. Blots of HA, GFP, and lamin from 1% of input material are shown. n=3. ADAR-HA and Zn72D-GFP interact in the presence of RNA. (G) Scatterplot of transcript enrichment in *Zn72D-GFP* head RIP-seq. Log_2_ fold change expression in RIP compared to 5% input is plotted versus the log of the average number of normalized counts of each transcript (dots) in input samples. Orange dots, transcripts have editing sites affected by Zn72D. The majority of transcripts with editing sites altered by Zn72D knockdown are enriched in the RIP. Black dots, *p* < 0.05, Wald tests. (H) Fold change of *qvr*, *cac*, *Shab*, and *para* transcripts recovered in *Zn72D-GFP* RIP compared to IgG negative controls, as measured by qPCR. Transcripts enriched in RIP as measured by RNA-seq (G) are also enriched when measured by qPCR.

We next tested whether ADAR and Zn72D proteins physically interacted. First, we immunoprecipitated ADAR-HA from *Adar-HA; Zn72D-GFP* fly head lysates and *Zn72D-GFP* fly head lysates as a negative control. Zn72D-GFP co-immunoprecipitated with ADAR-HA in the anti-HA IP in *Adar-HA; Zn72D-GFP* head lysates. However, after treatment with RNase A, the interaction was significantly weakened, suggesting that the two proteins interact in an RNA-dependent manner (**Figure 3E**). We subsequently performed the reciprocal co-IP in lysates from nuclei of heads of *Adar-HA; Zn72D-GFP* flies, using *Adar-HA* flies as a negative control. We found that ADAR-HA co-immunoprecipitated in the anti-GFP IP in nuclei containing both tagged proteins, but not after RNase A treatment (**Figure 3F**), which suggested that ADAR and Zn72D interact in an RNA-dependent manner within the nucleus.

As the RNA-dependent interaction between Zn72D and ADAR suggested that the proteins interact by binding the same RNAs, we hypothesized that Zn72D binds the transcripts with editing sites affected by *Zn72D* knockdown. We made multiple attempts to perform seCLIP-seq (Van Nostrand et al., 2017) on Zn72D-GFP from fly heads to determine Zn72D’s RNA binding sites, but saw little evidence of RNA binding, likely due to the inefficiency of crosslinking proteins to dsRNA *in vivo* using UV light (Wheeler et al., 2018). Instead, we performed RNA immunoprecipitation and sequencing (RIP-seq) on Zn72D by pulling down Zn72D-GFP and its bound RNAs from fly heads without crosslinking, extracting RNA from the inputs and IPs, and making RNA-seq libraries. For negative controls, we split lysates in half and incubated half with IgG antibody rather than GFP antibody; these negative controls did not immunoprecipitate enough RNA to amplify RNA-seq libraries, suggesting our pulldown was specific to RNAs bound by Zn72D-GFP. To determine transcript enrichment in the RIP, we counted the reads mapping to each gene in both the IP libraries and matched input libraries made from RNA extracted from 4% of the input lysates. We then used these counts as inputs to DESeq2 to determine genes with increased or decreased expression in the RIP compared to inputs. We found that of the 217 transcripts sequenced in the RIP with at least one editing site affected by Zn72D, 182 (84%) were significantly enriched in the RIP over the input (**Figure 3G, Table S4**). To further validate the results of the RIP-seq including our IgG negative controls, we used qPCR to quantify the relative levels of *qvr*, *cac*, *para*, and *Shab* in both IgG and GFP IPs and matched inputs (**Figure 3H**). We found that these transcripts with large editing changes in *Zn72D* knockdowns showed between 332- and 899-fold higher amounts in GFP IPs compared to IgG IPs after normalizing by input levels. Taken together, these experiments support the hypothesis that Zn72D binds at least some of the same transcripts that ADAR edits, which may help explain its role as a site-specific regulator of editing levels.

Since Zn72D appeared to bind many edited transcripts, and *Zn72D* and its human homolog *ZFR* both have reported roles in regulating pre-mRNA splicing (Haque et al., 2018; Worringer and Panning, 2007), we checked whether *Zn72D* knockdown led to splicing changes in the transcripts with editing changes. To identify alternative splicing changes in *Zn72D* knockdown brains (BDSC#55625, see **Figure S2**) compared to shGFP controls, we used MISO (Katz et al., 2010), which identifies differentially regulated isoforms across samples. We found that *Zn72D* knockdown altered splicing in 40 of the 252 transcripts where we observed editing changes (**Figure S4A**), and those 40 transcripts contain 216 of 785 editing sites (28%) altered by Zn72D. We found a total of 400 altered splicing events in 257 transcripts (**Figure S4B, Table S5**), suggesting Zn72D regulated both splicing and editing in a subset of transcripts and also regulated splicing and editing independently in some transcripts.

### Loss of Zn72D leads to impaired locomotion and neuromuscular junction defects

RNA editing is necessary for proper neuronal function in the fly (Jepson et al., 2011; Palladino et al., 2000), and loss of *Adar* leads to impaired locomotion and defects in neuromuscular junction (NMJ) morphology (Bhogal et al., 2011; Maldonado et al., 2013). Since loss of Zn72D led to such a dramatic change in RNA editing levels, we hypothesized that it might play a similar role to ADAR in regulating neuronal function. First, we tested locomotion in *Zn72D* knockdown flies. While Zn72D mutant flies die as pupae, *C155-Gal4; UAS-shZn72D* flies were viable into adulthood, allowing us to test their climbing ability using a negative geotaxis assay. We measured climbing in flies with *GFP* RNAi and *Zn72D* RNAi driven by *Elav-Gal4* by determining the proportion of flies of each genotype that climbed more than halfway up a 20 cm glass vial over time. We found that, while Zn72D knockdown flies can climb the sides of a glass vial, only an average of 36% of Zn72D RNAi flies climbed above 10 cm in a glass vial after two minutes compared to 100% of GFP RNAi flies (**Figure S5A**). This climbing defect was more severe than what we observed for *Adar* RNAi, suggesting that *Zn72D* knockdown lead to a locomotion phenotype that was distinct from *Adar* knockdown. In an independent test using *GFP* and *Zn72D* RNAi flies crossed to *C155-Gal4*, 46% of Zn72D RNAi flies were found above the 10 cm mark after 5 minutes, compared to 100% of GFP RNAi flies (**Figure S5B**).

To more deeply explore the cellular basis for this locomotor defect, we examined how loss of *Zn72D* affected the morphology and organization of synapses at the NMJ. ADAR is necessary for proper synaptic architecture and function at the NMJ (Bhogal et al., 2011; Maldonado et al., 2013), and as Zn72D regulates ADAR editing at many sites, we hypothesized that it may similarly be necessary for NMJ organization. In *Zn72D* mutants, we examined synaptic morphology and observed a 6-fold increase in the number of satellite boutons (**Figure 4A-D, I**), a defect typically associated with impaired endocytic cycling and BMP signaling (Dickman et al., 2006; O’Connor-Giles et al., 2008). In vesicle cycling mutants like *synaptotagmin I (syt I)*, *endophilin*, and *synaptojanin*, there is a marked increase in satellite bouton number. To determine whether any of these endocytic proteins were affected by the loss of Zn72D, we used immunocytochemistry to examine Syt I levels at the NMJ. At *Zn72D* mutant NMJs, Syt I levels are decreased by 31% (**Figure 4A-D, J**), suggesting a potential mechanism by which the loss of *Zn72D* results in excessive satellite boutons. Intriguingly, loss of ADAR increases levels of Syt I (Maldonado et al., 2013), suggesting that Zn72D and ADAR can regulate the levels of synaptic proteins differently. Consistent with this difference, *Adar* mutants lacked the increased number of satellite boutons (Bhogal et al., 2011), suggesting that both mutants regulate aspects of NMJ architecture differently. However, we also observed similarities between ADAR and Zn72D regulation of protein levels at the NMJ. Loss of *ADAR* also alters the levels of postsynaptic GluRIIA receptors (Maldonado et al., 2013); this is thought to be in response to changes in presynaptic function. Multiple allelic combinations of *Zn72D* mutants show a 32% reduction in synaptic GluRIIA staining (**Figure 4E-H, K**); this is consistent with the 37% reduction observed in *ADAR* mutants (Maldonado et al., 2013). Together with the morphology and Syt I staining, these results suggest that NMJ phenotypes arising from loss of *Zn72D* cannot be completely explained by a loss of ADAR editing. Rather, there are likely to be ADAR-dependent and ADAR-independent roles of Zn72D in regulating NMJ synapse organization. This hypothesis is consistent with the behavior of *Zn72D* knockdown flies in climbing assays and our observation that Zn72D regulated splicing in numerous transcripts where we did not see editing changes; Zn72D likely has functions inclusive of and beyond that of ADAR.

**Figure 4.**
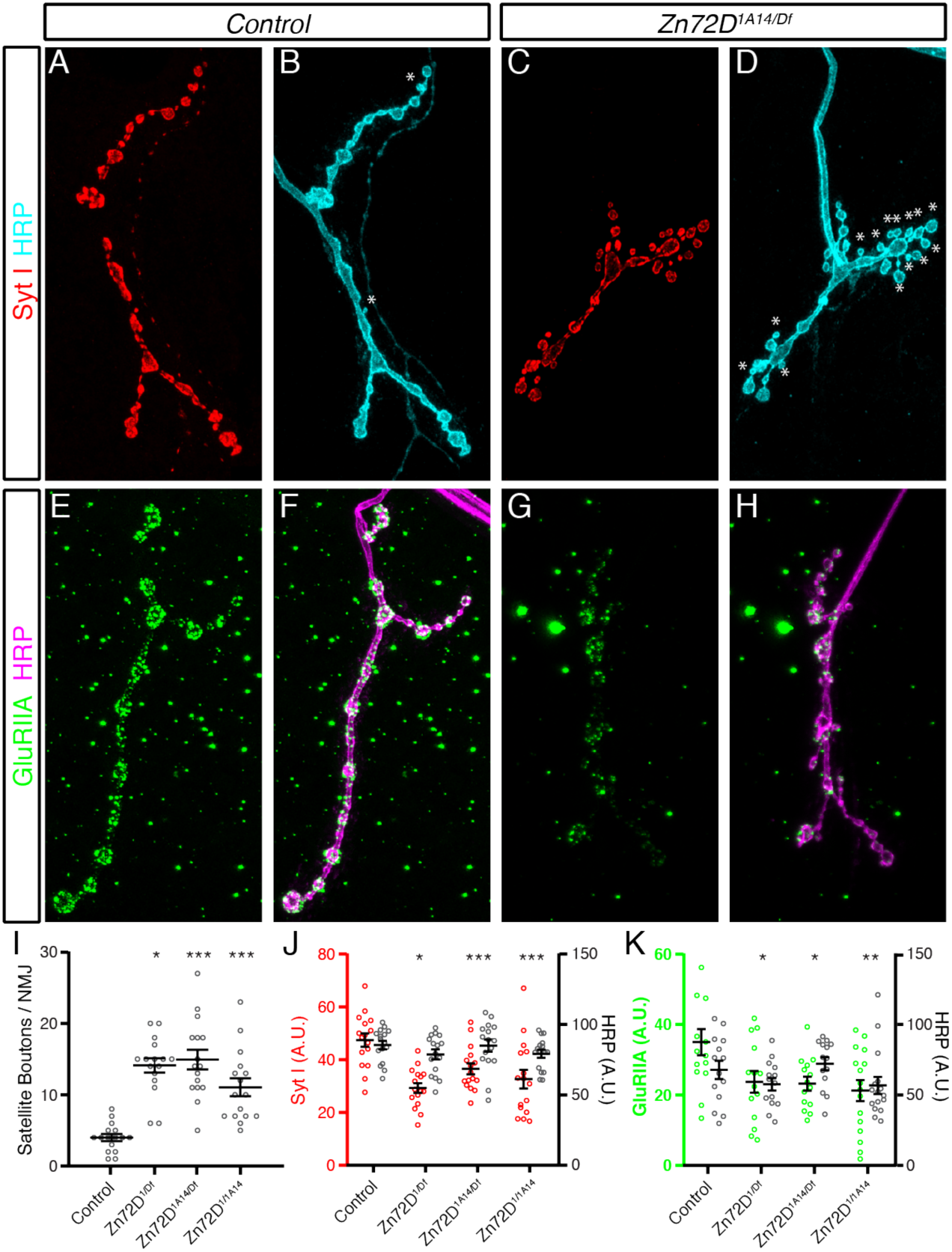
Loss of Zn72D Regulates NMJ Architecture and Protein Levels. (A-D) Third instar larvae stained with antibodies against Syt I (red) and HRP (blue) in Control (A-B) and *Zn72D^1A14/Df^* mutant larvae (C-D). Df is Df(3L)Exel6127, which lacks the *Zn72D* locus. Asterisks indicate satellite boutons. Loss of *Zn72D* markedly increases the incidence of satellite boutons. (E-H) Third instar larvae stained with antibodies against GluRIIA (green) and HRP (magenta) in Control (E-F) and *Zn72D^1A14/Df^* mutant larvae (G-H). Loss of *Zn72D* also results in reduced synaptic GluRIIA staining. (I-K) Quantification of satellite boutons per NMJ (I), Syt I fluorescence levels (J) and GluRIIA fluorescence levels (K). Multiple allelic combinations of *Zn72D* mutants show increased satellite bouton numbers and reduced Syt I and GluRIIA staining. HRP staining is unchanged across all genotypes, suggesting that these deficits are specific. For all graphs, open circles represent each individual value while the mean and S.E.M. are indicated by the error bars. In all cases, *n* ≥ 7 animals, 14 NMJs for each genotype. *, *p* < 0.05, **, *p* < 0.01, ***, *p* < 0.001. Statistics were determined using ANOVA followed by a Dunnett’s multiple comparisons test.

**Figure 5.**
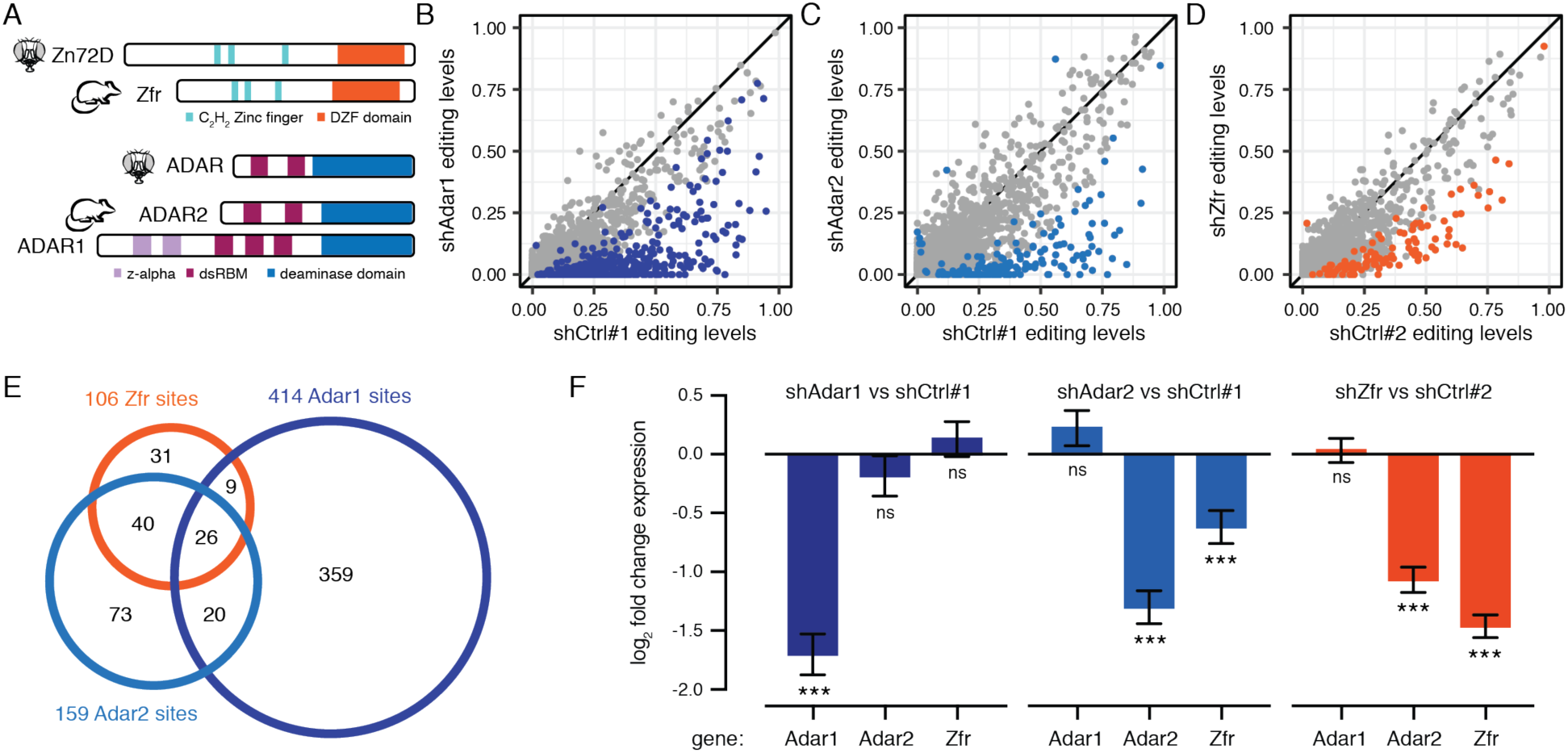
Zfr affects editing levels and *Adar2* mRNA levels in mouse primary neurons. (A) Schematic of protein domains of Zn72D and its mouse homolog, Zfr (top). Schematic of protein domains of dADAR and its mouse homolog ADAR2 along with the other catalytically active mouse ADAR, ADAR1 (bottom). (B) Comparison of editing levels between mouse primary neurons transfected with a control shRNA versus *shAdar1*. Blue dots, *p* < 0.05, Fisher’s exact tests. (C) Comparison of editing levels between mouse primary neurons transfected with control shRNA versus *shAdar2*. Blue dots, *p* < 0.05, Fisher’s exact tests. (D) Comparison of editing levels between mouse primary neurons transfected with control shRNA versus *shZfr*. Orange dots, *p* < 0.05, Fisher’s exact tests. Many editing sites show decreased editing upon *Adar1*, *Adar2*, and *Zfr* knockdown. (E) Venn diagram showing the overlap of affected sites between *shAdar1*, *shAdar2*, and *shZFR*. *shZfr* sites share a larger overlap with Adar2-affected sites, although the three sets are distinct. (F) Log_2_ fold changes of mRNA levels of *Adar1*, *Adar2*, and *Zfr* in *shAdar1*, *shAdar2*, and *shZfr* compared to *shControl* primary neurons. ***, *p* < 0.001, ns, *p* > 0.05, Wald tests. *Adar1* knockdown does not affect *Adar2* or *Zfr* levels, while *Adar2* knockdown decreases *Zfr* levels and *Zfr* knockdown decreases *Adar2* levels.

### Zn72D regulation of editing is conserved in mammalian neurons

We next wanted to determine whether the regulation of ADAR and RNA editing levels by Zn72D was conserved in mammals, as has been demonstrated for other regulators of RNA editing identified in flies (Bhogal et al., 2011; Hong et al., 2018b). To check whether the mammalian homolog of Zn72D, Zfr, altered editing levels in mammalian neurons, we designed shRNAs against mouse *Zfr* as well as *Adar2*, which encodes the homolog of dADAR, and *Adar1*, which encodes the other catalytically active mammalian ADAR protein (**Figure 4A**). We knocked down *Adar1*, *Adar2*, and *Zfr* in mouse primary cortical neurons, extracted RNA, and made and sequenced RNA-seq libraries. We compared editing levels between two combined biological replicates of primary neurons transfected with control shRNAs to those transfected with shRNAs against *Adar1*, *Adar2* and *Zfr* (**Figure 4B-D**). In each knockdown, we found more than 100 sites with decreased editing levels, demonstrating that *Zfr* knockdown alters editing levels in this mammalian neuronal context (**Table S6**). Among the sites affected by Zfr was the *Gria2* Q/R site that is known to play a critical role in neuronal function (Horsch et al., 2011). We compared the sites decreased upon knockdown of *Adar1*, *Adar2*, and *Zfr*, and we found all three knockdowns altered a distinct subset of editing sites. Of note, the set of sites decreased by *Zfr* knockdown more closely overlapped with those decreased by *Adar2* knockdown than those decreased by *Adar1* knockdown (**Figure 4E**). This finding is consistent with our findings in *Drosophila*, as the single ADAR enzyme is a closer homolog of mammalian ADAR2 in sequence and in function (Keegan et al., 2011).

Since Zfr appeared to affect mostly ADAR2-regulated sites, we used the RNA-seq data to measure *Adar1*, *Adar2*, and *Zfr* mRNA expression levels in all three knockdowns. We found that *Zfr* knockdown led to a decrease in *Adar2* mRNA expression (**Figure 4F**), suggesting that *Zfr* regulated *Adar2* levels in mouse primary neurons. Further supporting an ADAR2-centric role for editing level regulation by Zfr, we found that knocking down *ZFR* in human Hek293T cells (Haque et al., 2018) led to no change in editing (**Figure S6A-C, Table S7**). *Adar2* is much more lowly expressed in Hek293T cells than in the mouse primary neurons (**Figure S6D**), and therefore ADAR1 is likely responsible for the vast majority of editing events in these cells. Taken together, these data suggest that the broad mechanisms of Zn72D regulation of editing – regulating both ADAR levels and editing at specific sites – are conserved in the mouse brain between Zfr and ADAR2.

## Discussion

RNA editing is dynamically regulated during development and across tissue and cell types (Graveley et al., 2010; Sapiro et al., 2019; Tan et al., 2017; Wahlstedt et al., 2009), but few factors responsible for this regulation are known. Since RNA binding proteins (RBPs) like ADAR form extensive cross-regulatory networks (Dassi, 2017), they are top candidates for regulators of editing, and many of the known regulators of editing are proteins that interact with ADARs by binding the same RNAs (Quinones-Valdez et al., 2019; Washburn and Hundley, 2016). To identify additional RBPs that regulate editing, we performed an RNAi screen in the fly brain. Our previous work suggested a role for additional *trans* regulators of editing in the fly brain (Sapiro et al., 2019; 2015), and using *Drosophila* allowed us to perturb RBP levels *in vivo* rather than in cell lines to measure editing changes at a large number of editing sites through RNA-seq. The majority of the RNA binding proteins we screened had only a small influence on RNA editing levels at a few sites, suggesting that editing levels at a majority of sites are fairly stable even as the RNA binding protein landscape changes. This result is consistent with our previous findings that a large number of editing sites have stable editing levels across different neuronal populations in the fly brain (Sapiro et al., 2019), as well as a study of the role of RBPs in regulating editing levels in human cells (Quinones-Valdez et al., 2019). However, because we screened RNAi lines, many of which only reduced the targeted mRNA by around 50%, our screen results may include false negatives. Furthermore, future experiments probing double-stranded RNA binding proteins may lead to the identification of more critical *trans* regulators of editing, as ADARs interact with double-stranded RNA species (Bass, 2002).

Zn72D is a broadly influential, site-specific positive regulator of RNA editing, as a majority of editing sites in flies have reduced editing levels upon Zn72D knockdown (see **Figure 2**). To our knowledge, other than ADARs, Zn72D has the largest site-specific effect on regulating RNA editing levels, compared to previously identified regulators. Zn72D is a zinc finger RNA binding protein that was first identified as a suppressor of a mutation in the cell-cycle regulator *cyclin E* (Brumby et al., 2004). The Zn72D protein has three C_2_H_2_ zinc finger domains and a DZF domain that facilitates protein dimerization and contributes to RNA binding (Castello et al., 2016; Wolkowicz and Cook, 2012). The mouse homolog of Zn72D, Zfr, is predicted to bind A-form dsRNA helices due to similarities between its zinc finger domains and those of zinc finger proteins known to bind specifically to dsRNA: long linkers between zinc fingers, an interhistidine distance of five amino acids, and a reversal of characteristic aromatic and hydrophobic residues (Meagher et al., 1999). Zfr alters splicing in human macrophages, regulating innate immunity (Haque et al., 2018), which suggests it plays a broad role in RNA processing. Zn72D also regulates the male-specific lethal (MSL) dosage compensation complex in flies by altering the splicing of *maleless*, which encodes a critical member of the complex (Worringer and Panning, 2007). Interestingly, a gain-of-function mutation in *maleless* regulates RNA editing levels in *para* (Reenan et al., 2000), although loss-of-function mutations did not have the same effect. As the human homolog of Maleless, DHX9, is also known to regulate editing (Hong et al., 2018a), some of the Zn72D editing phenotype may be caused indirectly through regulation of *maleless*, although this possibility needs to be further explored in the future.

We found that *Zn72D* knockdown regulated editing in a site-specific manner, altering editing at the majority of sites but leaving a large number of editing sites unchanged. We showed that Zn72D co-localizes and interacts with ADAR in an RNA-dependent manner, leading us to hypothesize that Zn72D facilitates ADAR editing at some sites by binding the same dsRNAs as ADAR. While Zn72D enhances editing at a majority of editing sites, a large number of sites are unaffected by Zn72D levels and editing is inhibited by Zn72D at a small number of sites. The effect of Zn72D on editing differs within transcripts and even between sites found within a few bases of each other. While Zn72D loss leads to an overall decrease in ADAR protein, the effects of this loss are distributed asymmetrically across edited adenosines, in a manner that is distinct from the effect of knocking down *Adar* itself. While some of the observed editing decreases may be a consequence of lower ADAR levels, we hypothesize that for at least a subset of RNA species, the presence of Zn72D alters the efficiency at which particular adenosines are edited, specifically for the transcripts in which Zn72D loss increases editing.

The exact mechanics of the Zn72D-ADAR interaction need to be further studied, as Zn72D may affect editing in different ways. For example, Zn72D may alter the structure of ADAR-bound dsRNAs by modulating splicing kinetics; however, while we found 40 transcripts with both splicing and editing changes, there were many transcripts with editing changes where we did not find evidence of splicing changes. Furthermore, while splicing efficiency can alter editing levels (Licht et al., 2016), editing can also affect splicing (Hsiao et al., 2018), so future studies should determine whether Zn72D splicing is in fact regulating editing at some sites. Zn72D could also alter ADAR binding at certain dsRNA structures to change which adenosines get edited or modify ADAR’s ability to move along a substrate to edit multiple adenosines within a few bases of each other, a process which in itself is poorly understood. Future studies will help to clarify how Zn72D affects ADAR’s function.

In addition to molecular phenotypes, we found that loss of Zn72D leads to cellular and organismal phenotypes. *Zn72D* mutant larvae have abnormal morphology and protein expression at the NMJ, and *Zn72D* knockdown flies have decreased climbing ability, which are similar to defects found in *Adar* mutants. However, for both the NMJ and climbing defects, the phenotypes we observed differ somewhat from phenotypes that we or others have found in *Adar* mutants and knockdown flies. While some of the phenotypes we observed may stem from RNA editing defects, it is also likely that Zn72D has ADAR-independent functions. These results are also consistent with fact that *Zn72D* mutations cause lethality earlier than *Adar* mutations, and that Zn72D plays a role in regulating splicing outside of the transcripts where we observed editing changes. Overall, these results demonstrate that *Zn72D* plays a critical role in neurons, and that loss of Zn72D has important physiological consequences for the fly.

In mammalian neurons, we found that knocking down *Zfr* led to a large decrease in editing levels, suggesting that the neuronal role of Zn72D in RNA editing is conserved. Knockdown of *Zfr* affected mainly editing sites that were regulated by ADAR2, although at a subset distinct from those affected by *Adar2* knockdown. Knockdown of *Zfr* also led to a decrease in *Adar2* mRNA levels, suggesting that at least some portion of the editing phenotype may be due to decreased ADAR2 levels in mouse primary neurons. In a biochemical screen for proteins that interact with human ADARs, we identified human ZFR as a top ADAR1- and ADAR2-interacting protein and demonstrated an RNA-dependent interaction between ZFR and ADAR1 and ADAR2 (Freund et al., 2019). Together these results suggest that ZFR is also likely to be a direct site-specific editing regulator in mammals, although this mechanism could be further investigated through CLIP-seq experiments. Overall, in terms of both strength of editing effects across species and the breadth of sites affected within species, Zn72D/ZFR appears to be the most expansive regulator of RNA editing outside of the ADAR proteins discovered to date. The regulation of RNA editing by ZFR may have implications relevant to human disease. ADAR1 mutations can lead to Aicardi-Goutières syndrome (AGS) (Rice et al., 2012) and spastic paraplegia (Crow et al., 2014). These auto-immune diseases can have neurological symptoms and are caused by an increase in interferon expression accompanying a loss of ADAR1 editing of endogenous dsRNAs. ZFR has also been implicated, through identification of one missense mutation, in spastic paraplegia (Novarino et al., 2014). While our data suggest ZFR’s effect on editing is mainly exerted through ADAR2 rather than ADAR1, future studies should look to explore the consequences of this ZFR mutation on editing in more human contexts. Furthermore, as targeting ADAR1 has been shown to be an effective strategy to enhance cancer treatment (Ishizuka et al., 2018; Liu et al., 2018), ZFR – either through its regulation of editing or independent mechanisms of innate immune activation (Haque et al., 2018) – may prove to be a new candidate drug target. As a broadly influential *trans* regulator of RNA editing, detailed understanding of how Zn72D and ZFR regulate editing will provide novel insights into the editing process and how it can be disrupted.

## Methods

### Fly stocks and crosses

RNA binding protein shRNA lines for the screen were created as in (Ni et al., 2011); see **Table S1** for shRNA sequences and vectors used. *C155-GAL4* (BDSC#458) flies were obtained from Bloomington *Drosophila* Stock Center (BDSC), along with one *UAS-shAdar* line (BDSC#28311) and the independent *UAS-shZn72D* line (BDSC#55625) which were created by the Transgenic *Drosophila* RNAi project (TRiP) (Perkins et al., 2015). The stronger *shAdar* line was obtained from the Vienna *Drosophila* Resource Center (v7763) (Dietzl et al., 2007). For the RNAi screen, *C155-Gal4* virgins were crossed to males containing UAS-driven shRNAs against individual RNA binding proteins. If viable, 0-2 day old F1 females were collected and aged for three days. Approximately 15 brains were dissected from 3-5 day old females for each replicate, with two replicates per shRNA line. Zn72D-GFP (BDSC#50830) as well as Zn72D^1^ (BDSC#5061) and Zn72D^1A14^ (BDSC#32668) (Brumby et al., 2004), and Df(3L)Exel6127 (BDSC#7606) (Parks et al., 2004), which deletes chromosomal region 72D1-72D9 including *Zn72D* and surrounding genes, were obtained from BDSC. *Adar-HA^12.0.1^* (Jepson et al., 2011) flies were a generous gift from the R. Reenan lab and *Adar^5G1^* mutants (Palladino et al., 2000) a generous gift from L. Keegan. Flies were raised at 25°C on molasses-based food on a 12 hr light/dark cycle.

### RNA extraction and cDNA synthesis from fly heads and brains

RNA was extracted from dissected brains or heads using Agencourt RNAdvanced Tissue Kit (Beckman Coulter, Brea, CA: A32645) following the standard protocol but using one fourth of all volumes. To bind RNA to beads, final Bind Buffer was prepared by adding 10 ul of Bind Buffer beads to 90 ul of isopropanol. Following RNA extraction, 1 ul of TURBO DNase (Invitrogen, Carlsbad, CA: AM1907) was used to remove DNA by incubating for 20-30 minutes at 37°C. cDNA was synthesized from half of each RNA sample using SuperScript III (Invitrogen: 18080093) following the standard protocol using random hexamers as primers. The other half of the RNA was used as input for RNA-seq libraries.

### qPCR to test RNAi efficiency

qPCR was performed using KAPA SYBR Fast (Kapa Biosystems, Wilmington, MA: KK4600) to determine whether knockdown of the target exceeded 40% before proceeding to RNA-seq. qPCR primers were designed by FlyPrimerBank (Hu et al., 2013), and primer efficiency was tested to ensure 90-105% efficiency. qPCR was performed on a Bio-Rad CFX96 Real-Time System (Bio-Rad, Hercules, CA). Averaging three technical replicates, fold changes were calculated using the ΔΔCt method for the change between the gene of interest and reference gene *GAPDH*. Knockdown levels reported in Figure 1 were calculated using DESeq2 (Love et al., 2014) after RNA-sequencing.

### RNA-seq library preparation

rRNA was depleted from total RNA following RNase H-based protocols adopted from (Adiconis et al., 2013; Morlan et al., 2012). We mixed approximately 150 ng of RNA with 150 ng of pooled DNA oligos designed antisense to *Drosophila* rRNA in 50 bp sections (Supplemental Table 8) in an 8 ul reaction with 2 ul of 5X Hybridization buffer (500 mM Tris-HCl pH 7.4, 1 M NaCl). We annealed rRNA antisense oligos to total RNA samples for 2 minutes at 95°C, slowly reduced the temperature to 65°C and then added 2U of Hybridase Thermostable RNase H (Epicenter, Madison, WI: Lucigen H39500) and 1 ul of 10X Digestion buffer (500 mM Tris-HCl, 1 M NaCl, 200mM MgCl_2_) to make 10 ul total and incubated for 30 minutes at 65°C. rRNA-depleted RNA was then purified using 2.2X reaction volume of Agencourt RNAClean XP beads (Beckman Coulter: A63987), treated with TURBO DNase (Invitrogen: AM1907), and then purified with RNAClean XP beads again. rRNA-depleted RNA was used as input to KAPA Stranded RNA-seq Kit (Kapa Biosystems: KK8400) to make RNA-sequencing libraries for fly knockdowns. For mouse primary neuron RNA-seq libraries, the KAPA HyperPrep RNA-seq Kit (Kapa Biosystems: KK8540) was used to create libraries after rRNA depletion using oligos antisense to human rRNA sequences (Adiconis et al., 2013). All libraries were sequenced with 76 base-pair paired-end reads using an Illumina NextSeq (Illumina, San Diego, CA).

### Determining editing levels, gene expression, and splice junction usage from RNA-seq

RNA-seq reads were mapped using STAR v2.54b (--outFilterMultimapNmax 10 -- outFilterMultimapScoreRange 1 --outFilterScoreMin 10 --alignEndsType EndToEnd) (Dobin et al., 2013) to the dm6 genome (Aug 2014, BDGP Release 6 + ISO1 MT/dm6) (Hoskins et al., 2015). Mapped reads were filtered for primary hits only. Editing levels were determined using the Samtools v1.9 (H. Li et al., 2009) mpileup command to count A and G reads at known editing sites from (Duan et al., 2017; Graveley et al., 2010; Mazloomian and Meyer, 2015; Ramaswami et al., 2015; 2013; Rodriguez et al., 2012; Sapiro et al., 2015; St Laurent et al., 2013; Yu et al., 2016; Zhang et al., 2017). We required 20X coverage in each replicate, except for in *Zn72D* mutant versus wild type pupal head and mouse primary neuron comparisons, where we required 20X coverage total between the two replicates. Combined A and G counts from two replicates of each shRNA or mutant were compared combined A and G counts from two control replicates using Fisher’s exact test with a Benjamini-Hochberg multiple hypothesis testing correction in R v3.5.1 (Benjamini and Hochberg, 1995).

Gene expression levels were determined by counting reads hitting annotated genes in the transcriptome using RSEM v1.2.30 (B. Li and Dewey, 2011). RSEM outputted expected counts were rounded to the nearest integer and then used as input to DESeq2 (Love et al., 2014). The DESeq() and results() functions were used to calculate gene expression differences between pairs of cell types.

To analyze splicing changes in *Zn72D* knockdown flies, we trimmed all reads to 75 bp and then mapped reads using STAR v2.54b (--twoPassMode Basic), filtering for uniquely mapped reads. We ran MISO (Katz et al., 2010) after merging reads from two biological replicates of *shGFP* and *shZn72D* (BDSC#55625). We used the modENCODE *Drosophila* splice junctions available through MISO (https://miso.readthedocs.io/en/fastmiso/annotation.html), lifted over from dm3 to dm6 using UCSC Genome Browser LiftOver function (http://genome.ucsc.edu. After comparing events, we filtered for significant changes using --num-inc 1 --num-exc 1 --num-sum-inc-exc 10 --delta-psi .12 --bayes-factor 20.

### Brain immunofluorescence microscopy

3-5 day old female fly brains were dissected and stained exactly as in (Wu and Luo, 2006). The following primary antibodies were used: mouse anti-HA antibody (Covance, Burlington, NC: H11) and rabbit anti-GFP antibody (Abcam, Cambridge, UK: ab290) were used at 1:500, and rat anti-Elav antibody (Developmental Studies Hybridoma Bank, Iowa City, IA, deposited by G. M. Ruben: 7E8A10) was used at 1:25. Cross absorbed secondary antibodies used were: goat anti-mouse IgG Alexa Fluor Plus 555 (Invitrogen: A32727), goat anti-rabbit IgG Alexa Fluor 488 (Invitrogen: A11034), and goat anti-rat IgG Alexa Fluor 647 (Invitrogen: A21247). Brains were imaged on an Inverted Zeiss LSM 780 Multiphoton Laser Scanning Confocal Microscope (Carl Zeiss, Oberlochen, Germany) with a 20X objective.

### NMJ immunofluorescence microscopy

*Zn72D* mutant alleles were maintained over GFP-tagged balancer chromosomes or the larval-selectable Tb marker to enable selection as third instar larvae. Third instar larvae were dissected and stained as previously described (Mosca et al., 2012) in 0 mM Ca^2+^ modified *Drosophila* saline (Mosca et al., 2005). Larvae were fixed in 4% paraformaldehyde (Electron Microscopy Sciences, Hatfield, PA) for 20 minutes (for all antibodies except GluRIIA) or in Bouin’s Fixative (Electron Microscopy Sciences) for 5 minutes (for GluRIIA staining). The following primary antibodies were used: rabbit anti-Syt I at 1:4000 (Loewen et al., 2001), mouse anti-GluRIIA at 1:100 (Parnas et al., 2001), Cy5-conjugated goat anti-HRP at 1:100 (Jackson ImmunoResearch, West Grove, PA). The following secondary antibodies were used: Alexa-488 conjugated goat anti-mouse (Jackson ImmunoResearch) and Alexa568-conjugated goat anti-rabbit (Invitrogen), both at 1:250. Larvae were imaged on a Zeiss LSM 880 confocal microscope with a 40X, NA 1.3 or a 63X, NA 1.4 lens. NMJs on muscle 4 in segment A3 on both the right and left sides were imaged and quantified. All images were scored with the experimenter blind to genotype and processed using ImageJ (NIH) and Adobe Photoshop (Adobe, San Jose, CA). Immunofluorescence was quantified using ImageJ (NIH) and data analyzed using GraphPad Prism 8.0 (Graphpad Software, San Diego, CA).

### Co-immunoprecipitation experiments

Immunoprecipitation of HA-ADAR was performed as described in (Bhogal et al., 2011) with slight modifications as follows. Flies were flash frozen in liquid N_2_, their heads were removed by vortexing and then collected using a liquid N_2_ cooled sieve. Approximately 500 ul of fly heads were homogenized in lysis buffer (150 mM NaCl, 0.1% NP40, 20 mM HEPES (pH 7.4), 2 mM MgCl_2_, 1 mM DTT, cOmplete protease inhibitor (Sigma-Aldrich: 4693159001)) for input protein. Homogenates were centrifuged at 600 xg, supernatants were collected, and then additional lysis buffer was added, pellets were homogenized again, centrifuged, and then supernatants were collected and combined. Half of each lysate was treated with RNase A (Thermo Scientific, Waltham, MA: EN0531) for 30 minutes on ice. Equal amounts of lysate (approximately 1 mg) were rotated at 4°C overnight with 20 ul of mouse anti-HA agarose (Sigma-Aldrich, St. Louis, MO: A2095) and washed 5X for 10 minutes each with 1ml of lysis buffer. Protein was eluted in 2X Laemmli Sample Buffer (Bio-Rad: 161-0747) at 95°C for 10 minutes. Samples were run on 4-15% SDS-PAGE gels (Bio-Rad: 456-1086) and transferred to nitrocellulose membranes (Bio-Rad) for western blots. For immunoprecipitation of Zn72D-GFP, nuclei were collected from fly heads and immunoprecipitation was performed following the protocols described in (Piccolo et al., 2015), with slight modifications as follows. 20 ul of Protein G Dynabeads (Invitrogen: 10003D) were incubated with 5 ug of rabbit anti-GFP antibody (ab290). Following overnight incubation at 4C, IPs were washed 5 times with 1 mL of IP Wash Buffer, and Protein was eluted in 2X Laemmli Sample Buffer (Bio-Rad) at 95°C for 10 minutes. Samples were run on 4-12% SDS-PAGE gels (Bio-Rad) and transferred to nitrocellulose membranes (Bio-Rad) for western blots.

### Western Blotting

Antibodies used in western blots were: mouse anti-HA antibody (Covance: H11) at1:500, rabbit anti-GFP (Abcam: ab290) at 1:10000, mouse anti-GAPDH (Thermo Fisher: GA1R) at 1:2000, and mouse anti-lamin (Developmental Studies Hybridoma Bank, deposited by P. A. Fisher: ADL67.10-s) at 1:50 in 5% milk. Horseradish Peroxidase (HRP)-conjugated secondary antibodies (Jackson ImmunoResearch) were used 1:5000. Western blots were imaged after exposing to Pierce ECL Plus Western Blotting Substrate (Thermo Scientific: 32132) using a BioRad ChemiDoc imaging system running Image Lab Touch Software (v1.1.04). Quantification of western blots was performed using BioRad Image Lab 5.2. Bands were manually traced, and Adjusted Volumes of HA were normalized to GAPDH controls before comparisons between genotypes.

### RNA-immunoprecipitation and sequencing (RIP-seq)

RNA immunoprecipitation was performed after homogenizing ∼500 ul of fly heads in IP Buffer (150 mM NaCl, 20 mM HEPES pH 7.5, 2 mM MgCl2, 0.1% NP40, cOmplete protease inhibitor, RNaseOUT RNase inhibitor 1U/ul (Invitrogen)). Lysates were split in half, and 4% of input was removed for input control libraries. IP lysates were incubated overnight at 4C with Dynabeads Protein G (Invitrogen: 10003D), plus 5 ug of anti-GFP antibody (Abcam: ab290) or IgG (Sigma-Aldrich: I8765). IPs were washed 8 times in IP buffer. Beads and saved inputs were added to 1 mL of TRIzol (Thermo Fisher: 15596026). 200 ul of chloroform was added, and samples were centrifuged at 14000 xg at 4C for 15 minutes. Aqueous phases were collected, mixed with 1 volume of 70% ethanol and then transferred to a RNeasy MinElute column (Qiagen, Hilden, Germany: 74204) for purification following the standard protocol. RIP-seq libraries were made using KAPA HyperPrep RNA-seq Kits (Kapa Biosystems: KK8540) after rRNA depletion as described for RNA-seq library preparation. Libraries were sequenced with 76 base pair paired-end reads on an Illumina NextSeq. Reads were mapped to the dm6 as described above using STAR, and RSEM was used to obtain counts for reads hitting each gene. Expected read counts rounded to the nearest integer were used as input for differential expression analysis in DESeq2. Log2 fold changes were calculated using the DESeq() function followed by lfcShrink(type= “apeglm”) (Zhu et al., 2018). Normalized counts from inputs were determined using counts(normalized=TRUE). For qPCR, cDNA was made with iScript Advanced (Bio-Rad: 1708842), and qPCR was performed using KAPA SYBR Fast (Kapa Biosystems: KK4600) with 1 ul of input cDNA. GFP and IgG RIP Cts were normalized to inputs: ΔCt [RIP] = (Ct [RIP] – (Ct [Input] – Log2 (Input Dilution Factor))). Fold changes for each replicate were calculated as 2^−ΔCt[GFP] – ΔCt[IgG]^.

### Climbing assay

The negative geotaxis assay was performed with groups of 10 flies at a time counting the number of flies above the 10 cm mark on a glass vial every 30 seconds or 1 minute. Flies were given 24 hours to recover from CO_2_ exposure before tests.

### Lentivirus production

shRNAs targeted against *Adar1* (5’-CTCACTGAGGACAGGCTGGCGAGATGGTG), and *Adar2* (5’-AGCAATGGTCACTCCAAGTACCGCCTGAA), *Zfr* (5’-GAGTATACTGTGTTGCACCTTGGC), and non-targeting controls (control #1, matched with *Adar1* and *Adar2* knockdowns: 5’-ATCGCACTTAGTAATGATTGAA; control #2, matched with *Zfr* knockdown: 5’-AACCGATGTACTTCCCGTTAAT) were cloned into the pGreenPuro backbone from System Biosciences. This construct was used to produce a 6-well of lentivirus according to standard protocols in 293T cells using the third-generation system and concentrated 1:100 with lenti-X (Clontec). The virus pellet was stored at −80.

### Knockdown of ZFR, ADAR1, and ADAR2 in primary neurons

Primary mouse cortical neurons were dissociated into single cell suspensions from E16.5 mouse (strain: C57BL/6J) cortices using a papain dissociation system (Worthington Biochemical Corporation, Lakewood, NJ). Neurons were seeded onto poly-L-lysine coated plates (0.1% w/v) and grown in Neurobasal media (Gibco) supplemented with B-27 serum-free supplement (Gibco), GlutaMAX, and Penicillin-Streptomycin (Gibco) in a humidified incubator at 37°C, with 5% CO_2_. Half media changes were performed every 4-5 days, or as required. For gene silencing experiments, neurons were infected the day after seeding with a 6-well pellet worth of concentrated frozen virus (see above). The media was changed 12-16 hours later and every 4 days following (neurobasal + B-27 + glutamine). Neurons were harvested on day 7 and RNA was extracted using the PARIS kit from Ambion followed by TURBO DNase. Adar1, Adar2, and control shRNA#1 knockdowns were matched from the same mouse, while Zfr and control shRNA#2 knockdowns were matched from the same mouse. 250 ng of RNA was used to make next generation Illumina sequencing libraries using the KAPA HyperPrep kit from two biological replicates of each genotype. For *Adar2* knockdowns, we sequenced three technical replicates of the first biological replicate combined all A and G counts to increase coverage. Editing levels were determined after requiring 20X coverage total.

### Accession Numbers

The high-throughput sequencing data utilized in this work, including the RNA binding protein RNAi screen, Zn72D-GFP RIP-seq, and the mouse primary neuron RNA-seq, have been deposited in the Gene Expression Omnibus (GEO) database, accession number GSE126631.

## ACKNOWLEDGEMENTS

We thank members of the Li lab for their input throughout this project. We thank TRiP at Harvard Medical School (NIH/NIGMS R01-GM084947), The Bloomington Drosophila Stock Center (NIH P40OD018537), and the Vienna Drosophila Resource Center for fly stocks used in this study, and FlyBase for curating information from these resources. The Developmental Studies Hybridoma Bank, created by the NICHD of the NIH and maintained at The University of Iowa, Department of Biology, Iowa City, IA 52242, provided antibodies. Funding sources: NIH R01-GM102484 (to J.B.L.), R01-GM124215 (to J.B.L.), R01-MH115080 (to J.B.L.), National Science Foundation Graduate Research Fellowship DGE-114747 (to A.L.S.), NIH Training Grant NIH-NIGMS T32 GM007790 (to A.L.S.), American Heart Association Postdoctoral Grant 16POST27700036 (to E.F.), NIH R00-DC013059 (to T.J.M.), the Alfred P. Sloan Foundation (to T.J.M.).

## SUPPLEMENTARY FIGURES

**Figure S1.**
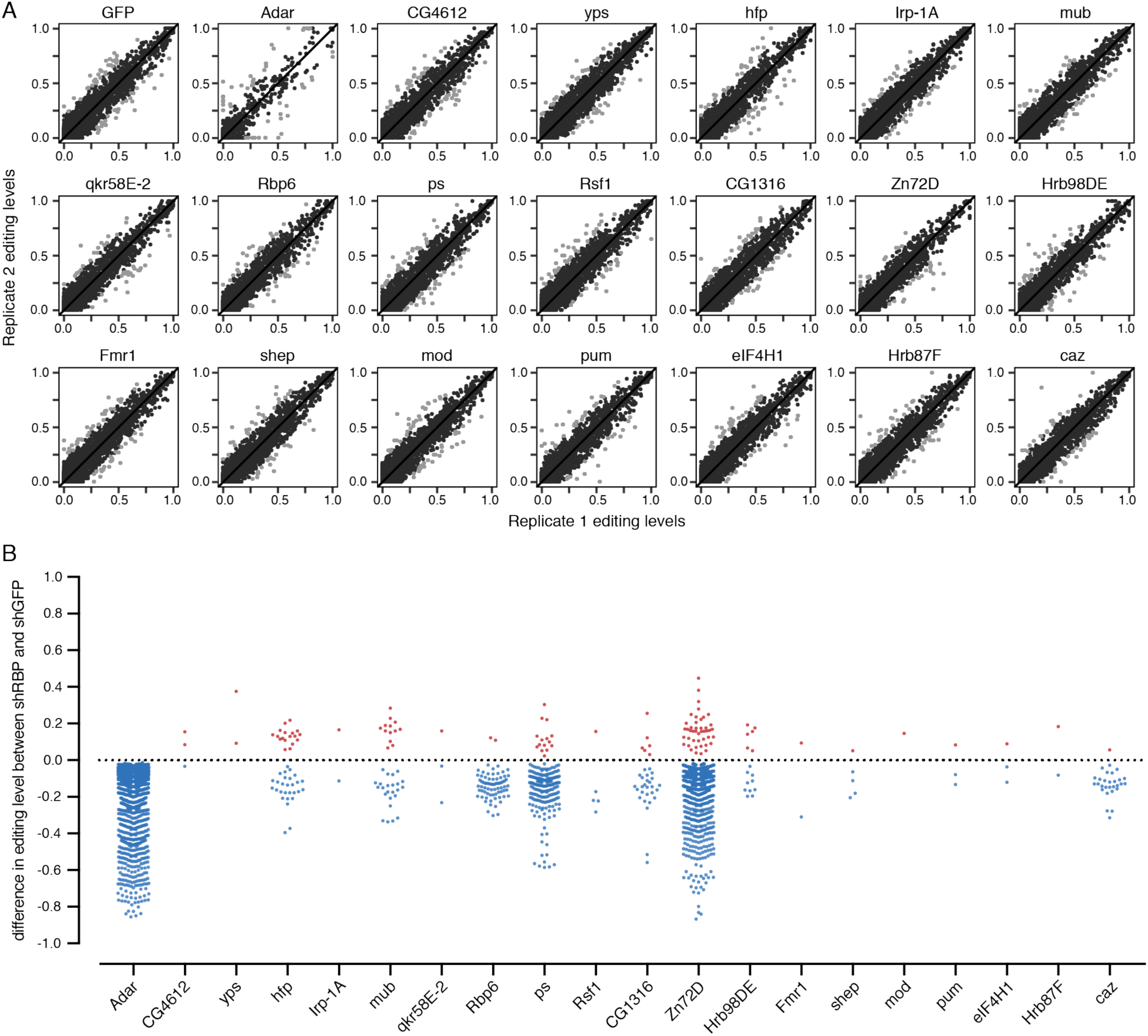
Biological replicates and editing level changes from RNAi screen. (A) Scatterplots of biological replicates from each knockdown used in the screen. All *UAS-shRBP* lines were crossed to *C155-Gal4*. Gray dots are more than 20% different and were not included in further analysis. All biological replicates showed highly reproducible editing levels. (B) The difference in editing between *shRBP* and *shGFP* for each significantly changed editing site in the RNAi screen. Blue and red dots, *p* < 0.05, Fisher’s exact tests. *shAdar* and *shZn72D* show the largest changes in editing.

**Figure S2.**
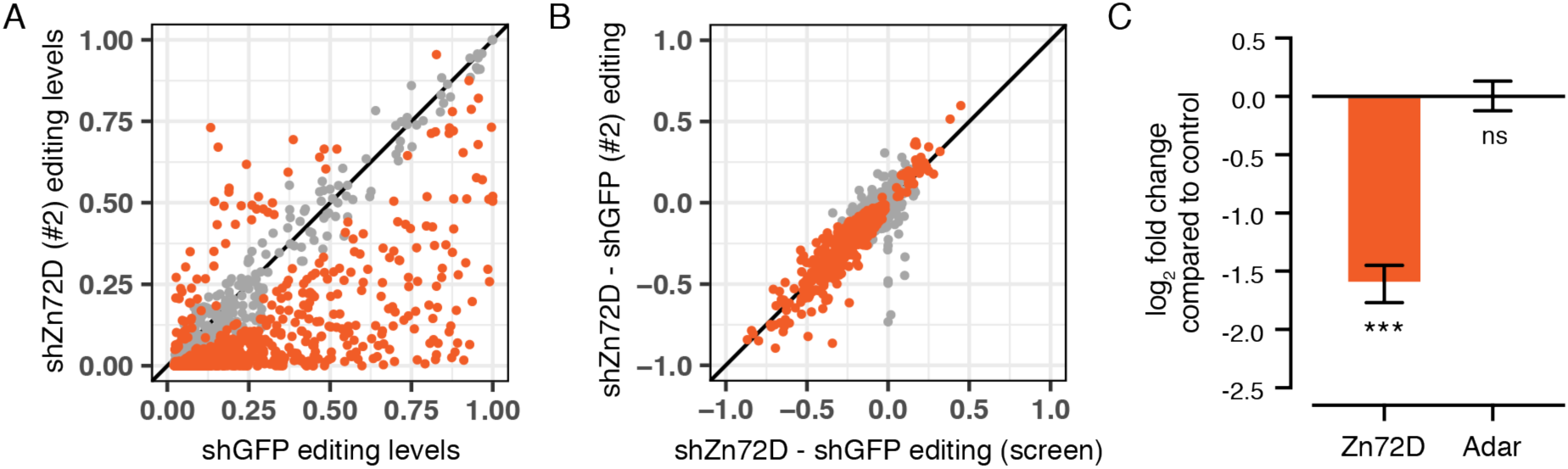
Independent shRNA knockdown confirms *Zn72D* RNAi editing phenotype. (A) Comparison of editing levels between *C155-Gal4; UAS-shGFP* (replicates 1 and 2) and *C155-Gal4; UAS-shZn72D* (shRNA#2: BDSC#55635). Orange sites, *p* < 0.05, Fisher’s exact tests. The majority of sites are altered by *Zn72D* knockdown. (B) Comparison of editing level differences between the two *shZn72D* lines and matched *shGFPs*. All lines were crossed to *C155-Gal4*. Orange sites *p* < 0.05, Fisher’s exact tests. The same editing sites are altered in each knockdown. (C) Log_2_ fold change of *Zn72D* and *Adar* mRNA levels in *shZn72D* (BDSC#55635) compared to *shGFP* control. ***, *p* < 0.0001, ns, *p* > 0.05, Wald tests. *Adar* mRNA levels are not changed in *Zn72D* knockdown.

**Figure S3.**
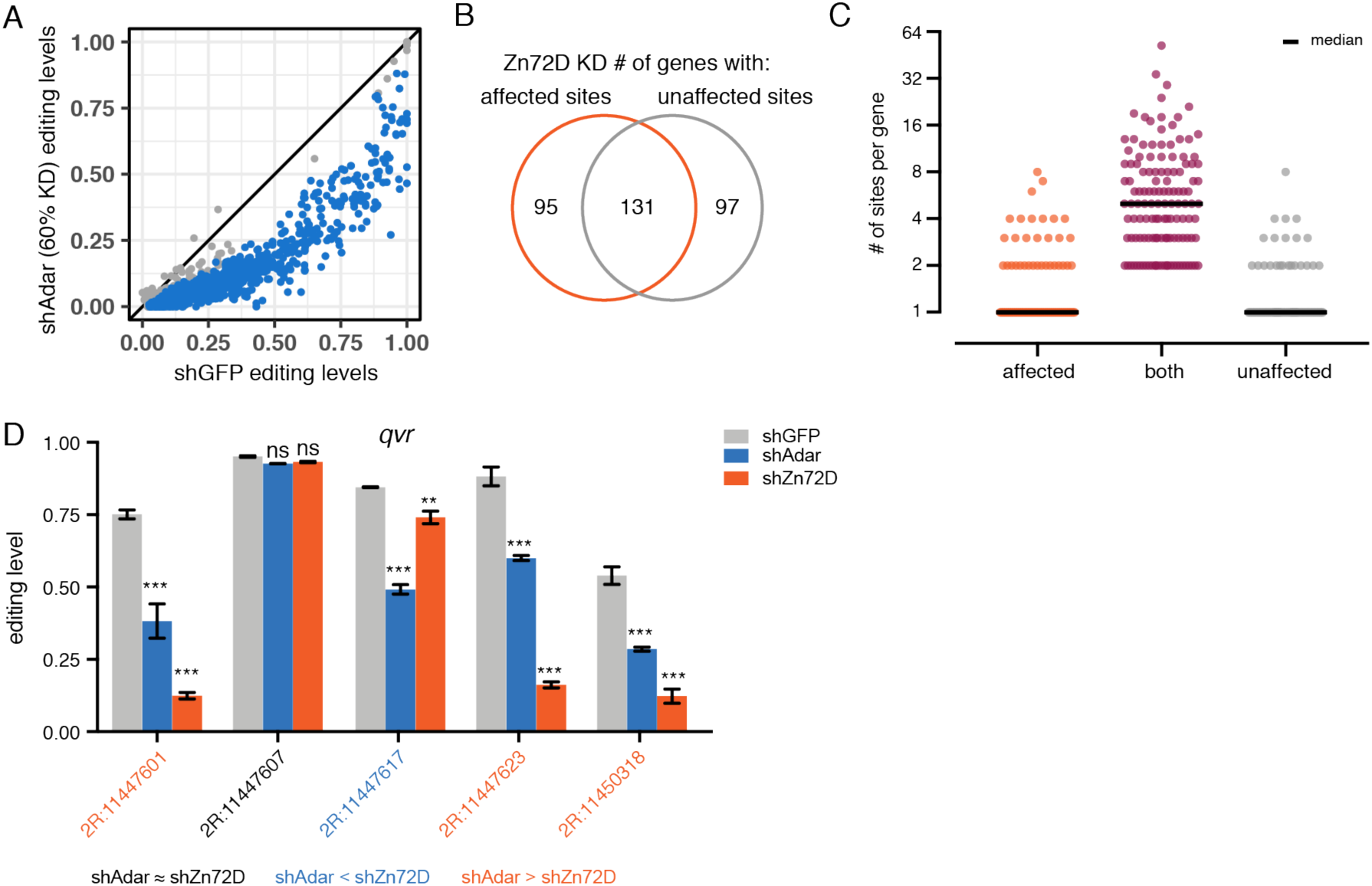
In contrast to *Adar* knockdown, *Zn72D* knockdown leads to varying changes at editing at sites found within the same transcript. (A) Comparison of RNA editing levels between *C155-Gal4; shGFP* and *C155-Gal4; shAdar* (60% decrease in *Adar* mRNA). Blue sites, *p* < 0.05, Fisher’s exact tests. The vast majority of sites are reduced. (B) The number of genes with editing sites that are either affected and/or unaffected by *Zn72D* knockdown. Some genes have only affected sites, some only unaffected, and many have both. (C) The number of editing sites per gene in genes with only affected, only unaffected or both affected and unaffected sites. Most transcripts with many editing sites show mixed effects of *Zn72D* knockdown. (D) Editing levels in *qvr* in *C155-Gal4; UAS-shGFP*, *C155-Gal4; UAS-shAdar* (60% KD), and *C155-Gal4; UAS-shZn72D*. Bars are mean and error bars are +/− SD, n=2. **, *p* < 0.01, ***, *p* < 0.0001, ns, *p* > 0.05. Asterisks over blue bars represent significance of change between *shGFP* and *shAdar*, and those over orange bars represent change between *shGFP* and *shZn72D*. The color of the coordinate represents the significance between *shAdar* and *shZn72D*, with black representing non-significant differences, blue representing *shAdar* levels lower than *shZn72D*, and orange representing *shZn72D* lower than *shAdar*, *p* < .001.

**Figure S4.**
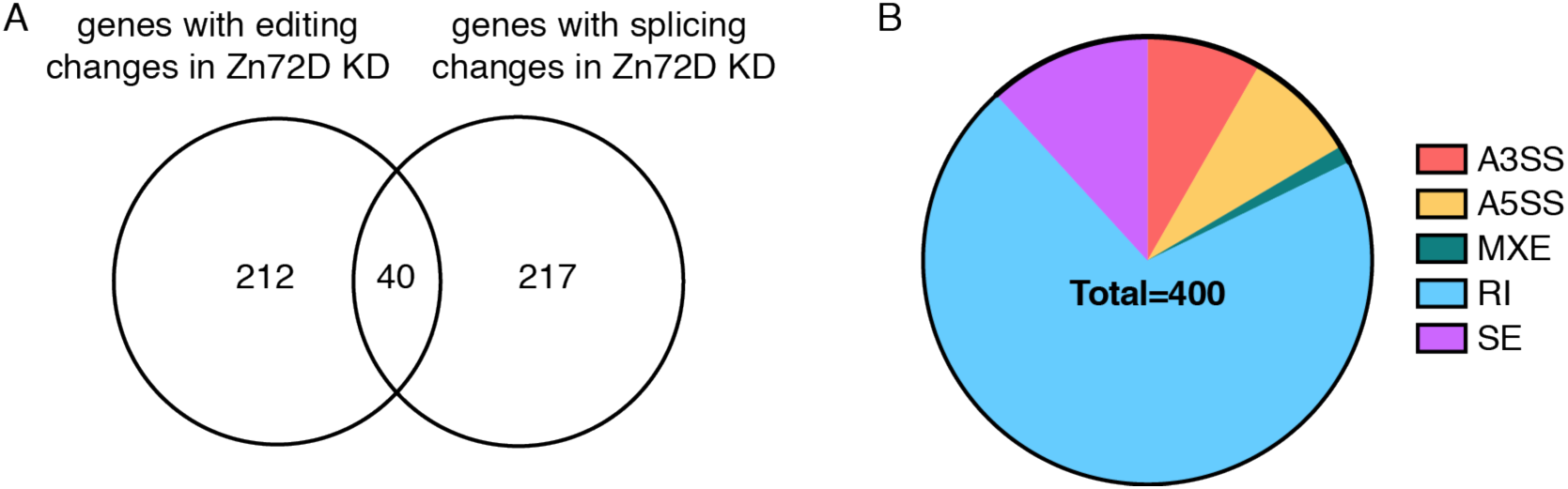
Overlap of genes with Zn72D-altered editing and splicing. (A) The number of genes with affected editing and/or splicing in *C155-Gal4; shZn72D* (shRNA#2: BDSC#55635) compared to *C155-Gal4; shGFP*. (B) The total number of splicing changes caused by *Zn72D* KD, split into type of splicing event. A3SS = alternative 3’ splice site, A5SS = alternative 5’ splice site, MXE = mutually exclusive exons, RI = retained intron, SE = skipped exon.

**Figure S5.**
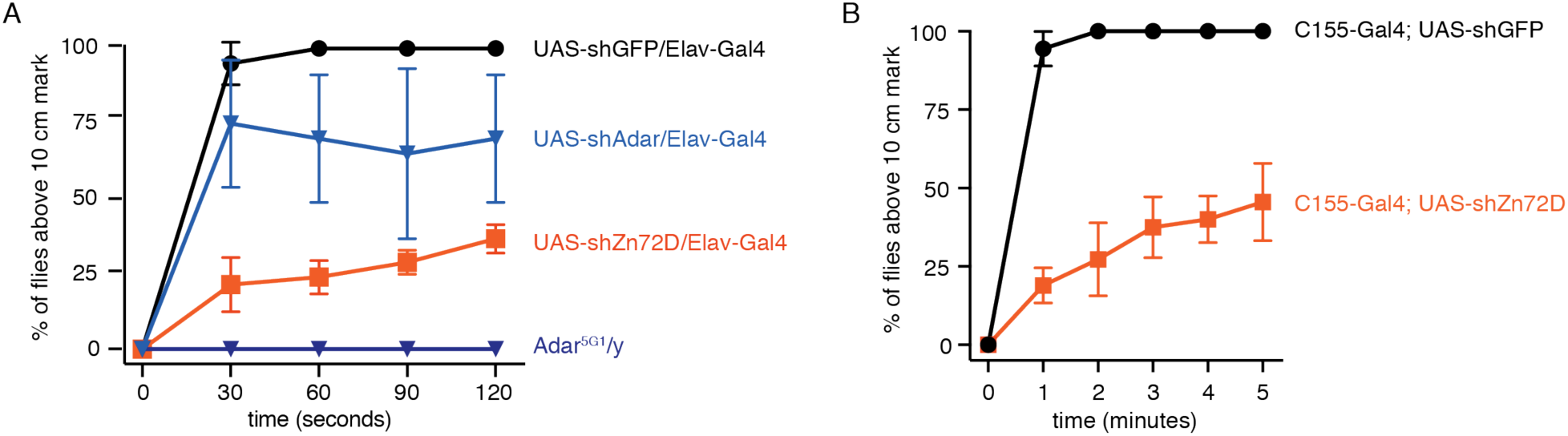
*Zn72D* knockdown causes a climbing defect in adult flies. (A) Negative geotaxis assay measuring the percentage of flies to reach the top half of a 20 cm glass vial over two minutes. shRNA lines were crossed to *Elav-Gal4*. n=4 sets of 10 flies. *Zn72D* knockdown has a more severe climbing defect than *Adar* knockdown, but not than *Adar^5G1^* mutants. (B) Negative geotaxis assay measuring the percentage of flies to reach the top half of a 20 cm glass vial over five minutes. shRNA lines were crossed to *C155-Gal4*. n=4 sets of 10 flies. The Zn72D knockdown climbing phenotype is reproducible and is maintained after five minutes.

**Figure S6.**
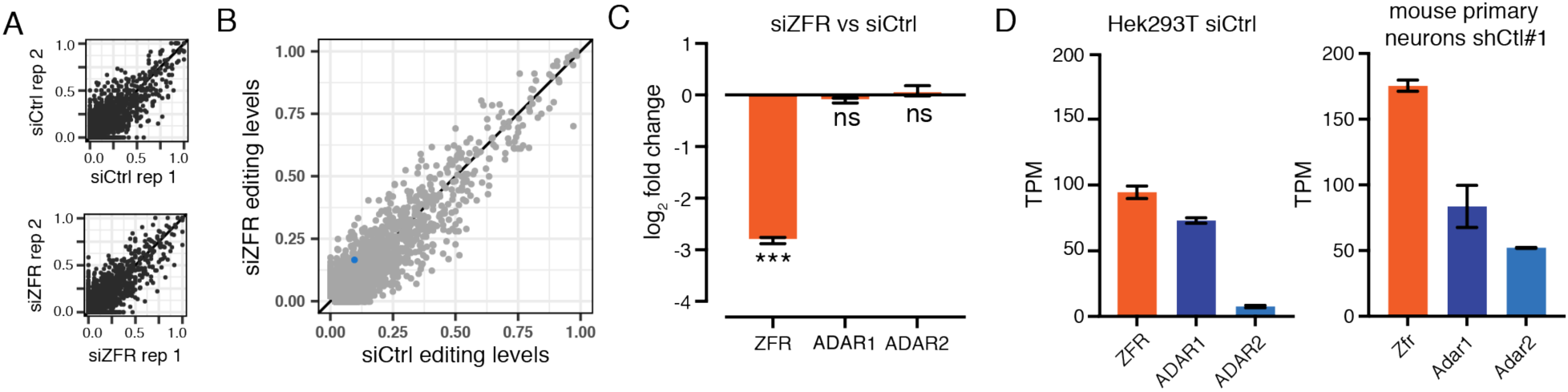
*Zfr* knockdown does not affect editing levels in Hek293T cells. (A) Representative comparisons of editing levels between biological replicates of *siCtrl* and *siZFR* in Hek293T cells (Haque et al. 2018). Editing levels are highly reproducible. (B) Comparison of editing levels between *siCtrl* and *siZFR*. Blue site, *p* < 0.05, Fisher’s exact tests. ZFR knockdown does not alter editing in Hek293T cells. (C) Log_2_ fold changes of *ZFR*, *ADAR1* and *ADAR2* in *siZFR* compared to *siCtrl*. ***, *p* < 0.001, ns, *p* > 0.05, Wald tests. (D) Transcripts per kilobase million (TPM) of *ZFR*, *ADAR1*, and *ADAR2* in Hek293T cells (left) and *Zfr*, *Adar1*, and *Adar2* in mouse primary neurons. *Adar2* is very lowly expressed in Hek293T cells in contrast to mouse primary neurons.

## LIST OF SUPPLEMENTARY TABLES

**Supplementary Table 1. UAS-shRNA fly lines used in this study.**

**Supplementary Table 2. Fly brain RNA binding protein knockdown editing levels and p-values from Fisher’s exact tests in comparisons to GFP RNAi.**

**Supplementary Table 3. Log_2_ fold changes and p-values of shRNA target genes and Adar in RBP knockdowns.**

**Supplementary Table 4. RNA-immunoprecipitation enrichment fold changes and p-values.**

**Supplementary Table 5. Zn72D-affected splice junctions as determined by MISO.**

**Supplementary Table 6. Mouse primary neuron editing levels for Zfr, Adar1, and Adar2 knockdowns and p-values from Fisher’s exact tests in comparisons to controls.**

**Supplementary Table 7. HEK293T editing levels for ZFR knockdown and p-values for comparison with control.**

**Supplementary Table 8. Drosophila rRNA antisense oligos used for rRNA depletion.**

